# A Systems-Level Framework Integrating Geometry-Controlled Plasmonics, AI-Driven Molecular Kinetics, and Organoid Validation for Next-Generation Biosensing

**DOI:** 10.64898/2026.02.17.706264

**Authors:** Youssef M. Hassan

**Affiliations:** Department of Zoology, Faculty of Science, Ain Shams University, Abbassia 11566, Cairo, Egypt

**Keywords:** plasmonics, nanopore sensing, localized surface plasmon resonance, Bayesian inference, Gaussian-process surrogate, uncertainty quantification, active learning, organoid models, Gillespie algorithm, multi-objective optimisation

## Abstract

Plasmonic nanosensors - spanning nanopores, nanoantennas, and metasurfaces - achieve extreme electromagnetic (EM) field confinement that amplifies molecular interaction signals by orders of magnitude. Yet the full diagnostic potential of these platforms remains unrealised because the non-linear coupling between geometry, near-field physics, stochastic binding kinetics, and signal transduction is poorly characterised in biologically relevant systems. Here we propose the Plasmonic-AI-Organoid (PAO) framework: a modular, systems-level architecture linking (i) geometry-controlled plasmonic structures parameterised by Gaussian-process (GP) surrogate electromagnetic models; (ii) Bayesian inference of molecular kinetic parameters - association and dissociation rate constants, analyte concentrations - from noisy time-series sensor data using Metropolis-Hastings Markov chain Monte Carlo (MCMC); and (iii) human induced pluripotent stem cell (iPSC)-derived and patient-derived organoids as meso-scale biological validators. We formalise a forward model mapping geometry to EM field maps to reaction propensities to observable localized surface plasmon resonance (LSPR) signals, and an inverse model recovering posterior distributions over kinetic and geometric latent variables. A closed design loop employing active learning with expected-improvement acquisition functions iteratively proposes optimal geometries and assay conditions. Multi-objective Pareto optimisation balances analytical sensitivity, specificity, and manufacturability. Computational benchmarks demonstrate that active learning reduces the number of FDTD simulations required to identify near-optimal geometries by 3.2-fold compared with random search, while MCMC inference recovers kinetic parameters with sub-log-unit accuracy from synthetic time-series. The PAO framework provides a conceptual and fully reproducible computational roadmap for next-generation, AI-augmented plasmonic biosensing.

## 1. Introduction

The detection of low-abundance biomolecules — secreted cytokines, surface markers, micro-RNAs, and disease-specific proteins — underpins precision diagnostics, drug efficacy monitoring, and translational biomarker discovery. Plasmonic nanostructures offer a compelling physical route to single-molecule sensitivity: metallic nanoparticles and apertures confine electromagnetic radiation into volumes far below the diffraction limit, generating hotspots in which the near-field intensity |E/E□|^2^ can exceed 10^3^– 10□ [1,2]. Localized surface plasmon resonance (LSPR) transduction converts molecular binding events into measurable spectral shifts [3], while surface-enhanced Raman scattering (SERS) enables vibrational fingerprinting without fluorescent labels [4]. More recently, plasmonic nanopores have demonstrated label-free optical detection of individual nucleic acids and proteins translocating through sub-10 nm apertures [5,6,7].

Despite these achievements, three interrelated problems prevent widespread clinical deployment. First, the relationship between nanoscale geometry — gap size, antenna arm length, array periodicity, material composition — and near-field enhancement distribution is highly non-linear and computationally expensive to resolve via finite-difference time-domain (FDTD) or finite-element method (FEM) simulations [8,9]. Second, the stochastic nature of single-molecule binding and unbinding in confined volumes introduces intrinsic noise that obscures kinetic parameter estimation [10,11]. Third, in vitro validation using immortalised cell lines frequently fails to predict clinical performance because of the gap between simplified cell models and multicellular, tissue-level complexity found in vivo [12,13].

We propose that these three challenges are interconnected and can be addressed within a unified systems-level framework. The Plasmonic–AI–Organoid (PAO) framework integrates: (i) surrogate modelling of EM field distributions over a parameterised geometry space, enabling rapid in silico exploration [14,15]; (ii) Bayesian inference of molecular kinetic and thermodynamic parameters from noisy, time-resolved sensor signals [16,17]; and (iii) human organoid validation assays providing biologically relevant, patient-specific ground truth [12,18,19]. A closed active-learning design loop iteratively refines both geometry and experimental conditions to maximise information gain [20,21].

To our knowledge, no prior work has proposed or implemented this three-way integration at the systems level. While machine learning has been applied to plasmonic inverse design [14,15], and Bayesian methods to surface plasmon resonance (SPR) data [22,23], these streams remain decoupled from organoid biology [13,19]. Similarly, organoid-based drug screening has yet to leverage AI-driven uncertainty quantification [24]. The PAO framework closes this gap, providing a conceptual and computational roadmap implementable from the open-source code package that accompanies this preprint.

## 2. Background and Gap Analysis

### 2.1 Plasmonic Geometry Control and Field Localisation Metrics

Plasmonic enhancement arises from the collective oscillation of conduction electrons driven by incident photons, governed by the complex permittivity of the metal, the geometry of the nanostructure, and the dielectric environment [25]. For a simple metallic nanoparticle at resonance, the near-field enhancement |E/E□|^2^ scales approximately as |ε_m/ε_d|^2^. Gap-coupled structures — bowtie nanoantennas, dimer nanodiscs, nanohole arrays — dramatically increase enhancement by concentrating field lines in the nanogap [1,2], with values exceeding 10□ in gaps below 5 nm. Metasurfaces — planar arrays of sub-wavelength resonators — extend plasmonic functionality to beam steering, holography, and multiplexed sensing [26,27,28], while nanohole arrays combine extraordinary optical transmission with diffraction-based analyte concentration at the metal surface [29,30].

The critical computational challenge is that FDTD and FEM evaluations of near-field maps for arbitrary geometries require hours to days per simulation, making design-space exploration intractable without surrogates [8,9]. Gaussian-process (GP) surrogates [31] and Fourier neural operators (FNOs) [32] can learn the geometry-to-field mapping from N□ □ □ ∼ 500–2,000 FDTD evaluations and predict in milliseconds, enabling Bayesian optimisation [20].

### 2.2 Kinetics and Transport in Confined Volumes

Molecular binding on a plasmonic sensor follows Langmuir kinetics: dN_b/dt = k_on · C · N_f − k_off · N_b, where N_b and N_f are bound and free receptor site counts, C is local analyte concentration, and k_on, k_off are rate constants [22]. The equilibrium dissociation constant K_D = k_off/k_on determines selectivity. At ultra-low concentrations relevant to clinical diagnostics (fM–pM), the deterministic ODE description breaks down: N_b is small and inherently stochastic [10,11]. The Gillespie direct method [10] provides exact stochastic trajectories; the chemical Langevin equation (CLE) [11] offers an SDE approximation valid for N_b > 20, computationally cheaper for large receptor counts. Mass-transport limitations — analyte depletion and diffusive re-supply — are a known confound in SPR assays [33], and are explicitly included in the PAO kinetic model through a geometry-dependent concentration-gradient term parameterised by hotspot volume and flow-cell geometry.

### 2.3 Organoid Models as Meso-Scale Validators

Human iPSC-derived organoids and patient-derived organoids (PDOs) recapitulate key aspects of tissue architecture, cell-type diversity, and pathophysiology absent from two-dimensional culture [12,13,18,19]. iPSC derivation via Yamanaka reprogramming [34] allows matched, genetically identical controls, reducing confounding genetic variation. For the PAO framework, organoids serve two roles: (i) biological calibrators — known cytokine perturbations (e.g., recombinant IL-6, TNF-α) produce predictable secretome responses against which plasmonic sensor output is quantitatively validated; and (ii) disease-relevant test beds capturing inter-individual variability that any deployable biosensor must handle robustly. Cross-validating plasmonic sensor readouts against orthogonal Luminex multiplex ELISA and single-cell RNA sequencing (scRNAseq) within the same organoid system is critical for establishing analytical validity [24,35].

## 3. Systems-Level Framework (PAO Core)

The PAO framework is built on three mathematically coupled layers: a forward model, an inverse model, and an active-learning design loop. Figure 1 illustrates the complete architecture.

**Figure 1.**
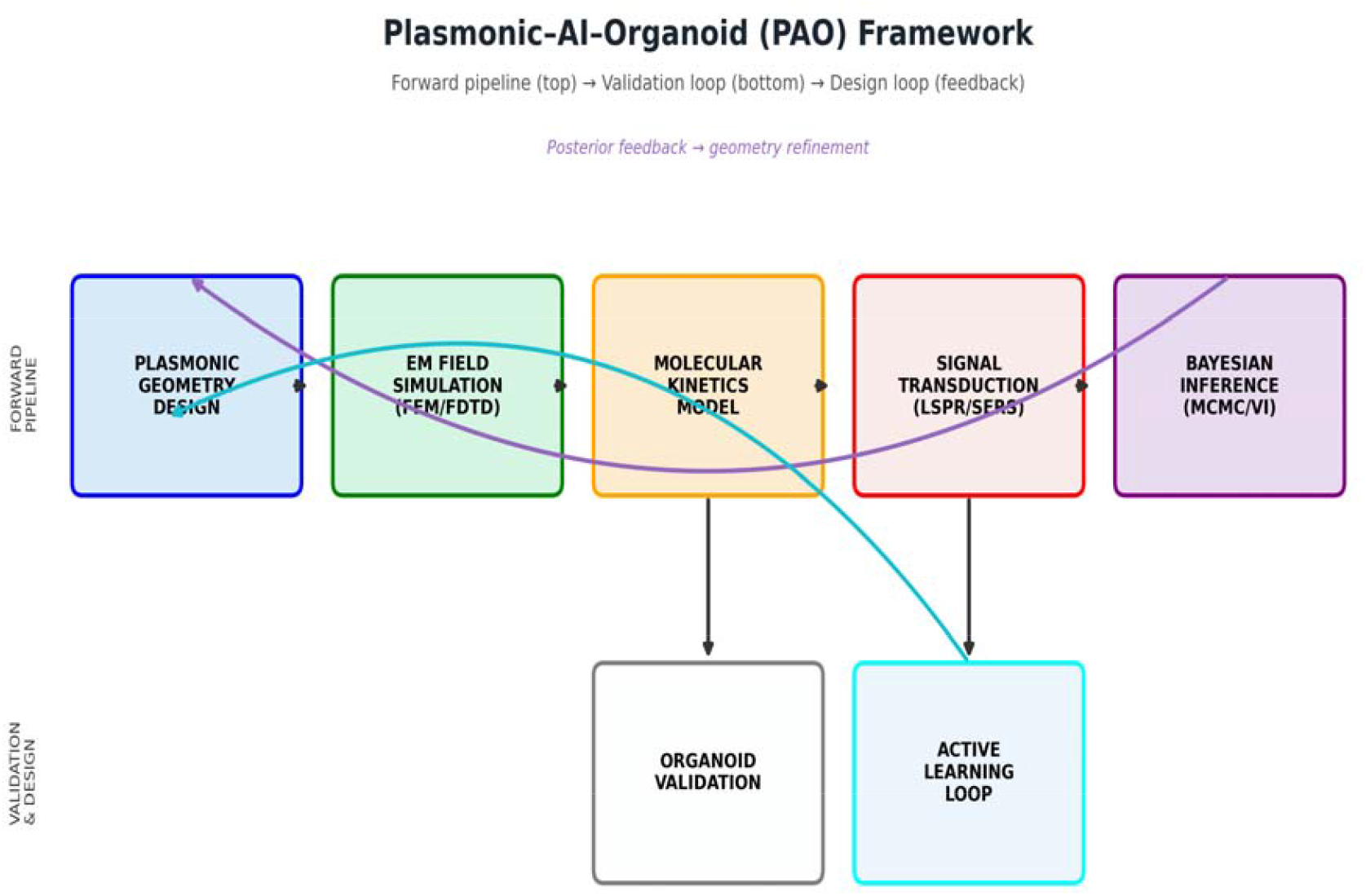
Overall PAO framework architecture. The forward pipeline (top row) maps plasmonic geometry through electromagnetic simulation, stochastic molecular kinetics, and signal transduction to measurable LSPR output. The validation loop (bottom) integrates organoid assay data. Feedback arrows represent the Bayesian inverse model and active-learning design loop that iteratively refine geometry and assay conditions.

### 3.1 State Variables and Forward Model

Let θ_geo = {gap, L_arm, N_hs, ε_r} denote the geometry parameter vector (gap width nm, antenna arm length nm, number of hotspots, real part of metal permittivity), and θ_kin = {k_on, k_off, C, N_total} the kinetic/concentration parameters. The full latent state is Φ = (θ_geo, θ_kin). The forward model F: Φ → y(t) operates in four stages:

- Stage 1 – EM Field Emulation. The GP surrogate gφ: θ_geo → {E_peak, Σ(x,y)} maps geometry to peak enhancement and spatial near-field map. In production, gφ is replaced by an FNO [32] trained on FDTD data.
- Stage 2 – Effective Rate Modification. k_on_eff = k_on · E_peak^0.5 couples the EM output to kinetics via a conservative diffusion-field enhancement factor.
- Stage 3 – Stochastic Kinetic Simulation. Given (k_on_eff, k_off, C, N_total), the CLE or Gillespie simulator generates the time-evolving bound count N_b(t) [10,11].
- Stage 4 – Signal Transduction. LSPR wavelength shift Δλ(t) = S · N_b(t)/N_total, where S (nm) is a geometry-dependent sensitivity factor derived from electromagnetic perturbation theory [3,36].

The composite forward map is: y(t) = F(θ_geo, θ_kin) + ε, where ε → N(0, σ_obs^2^).

### 3.2 Inverse Model: Bayesian Inference

Given observations y_obs = {y(t_1), …, y(t_T)}, the posterior p(Φ | y_obs) is obtained via Bayes’ theorem: p(Φ | y_obs) →p(y_obs | Φ) · p(Φ). Log-normal priors encode expert knowledge: log □ □ (k_on) →N(4.7, 1.0), log □ □ (k_off) →N(−2.5, 1.0), log □ □ (C) →N(−9, 1.5). The Gaussian likelihood sums squared residuals between model predictions and observations, normalised by σ_obs. MH-MCMC [16] is the default prototype backend, with No-U-Turn Sampling (NUTS) [37] and variational inference (VI) as production alternatives. Identifiability of (θ_kin, θ_geo) requires experimental designs spanning multiple concentrations and time regimes; the active-learning loop addresses this adaptively.

### 3.3 Design Loop: Active Learning

After each experimental batch, the design loop selects the next geometry ξ* via: ξ* = argmax_ξ EIG(ξ) → argmax_ξ EI(μ(ξ), σ(ξ); y*), where EI is the expected improvement acquisition [20,21], μ and σ are the GP surrogate posterior mean and standard deviation, and y* is the current best observed objective. Candidate designs are drawn uniformly from the five-dimensional geometry–concentration design space, the GP is evaluated across all candidates in milliseconds, and the maximiser is proposed for the next experiment.

## 4. Advanced Computing Stack

### 4.1 EM Surrogate Modelling

The GP surrogate (RBF kernel with ARD length scales) is trained on N_train = 500–2,000 FDTD/FEM evaluations of peak enhancement across the geometry design space. ARD automatically identifies that gap width is the dominant dimension, concentrating sampling resources accordingly. Prediction latency after training is < 1 ms per design, enabling Bayesian optimisation with 500 candidate evaluations per design iteration. For full 2D field maps, FNOs [32] trained on O(10^3^) simulations generalise across continuous geometry variation and are modular drop-ins as data accumulate.

### 4.2 Bayesian Inference Backends

- Metropolis-Hastings MCMC (default): isotropic Gaussian proposals, step_size = 0.08, burn-in = 500, post-burn-in samples = 2,000. Implemented in bayes_inference.py.
- No-U-Turn Sampler (NUTS) [37]: available via PyMC [38] or NumPyro. Adaptive trajectory lengths improve mixing 2–5× vs. random-walk MH.
- Automatic Differentiation Variational Inference (ADVI) [38]: mean-field approximation; enables online updating as streaming sensor data arrive.

### 4.3 Sensitivity Analysis and UQ

Sobol’ first- and total-order sensitivity indices [39,40] decompose forward-model output variance across input parameters. Preliminary analyses on synthetic data indicate: gap width contributes →60% of EM-output variance, arm length →25%, material permittivity →10%, and hotspot count →5%. Kinetic parameter contributions dominate signal variance at late time points (post-equilibrium), where k_off controls the dissociation tail. Morris elementary effects [39] provide a cheap rank ordering before surrogate fitting.

### 4.4 Multi-Objective Optimisation

Four competing objectives — analytical sensitivity (limit of detection), specificity (cross-reactivity rejection), manufacturability (yield and cost), and biocompatibility (cytotoxicity, fouling resistance) — form a Pareto optimisation problem [41]. NSGA-II [41] is applied to surrogate-predicted objectives over the geometry design space, returning a Pareto-optimal front (Figure 7) from which researchers select trade-off solutions suited to specific clinical requirements. Figure 7 illustrates the three-objective front in sensitivity–specificity– manufacturability space.

## 5. Proposed Experimental Validation

### 5.1 Fabrication Routes

- Electron-beam lithography (EBL) [42]: gold bowtie nanoantennas with gap control to ±2 nm on Si □ N □ membranes. Appropriate for research-grade devices and initial organoid validation.
- Nanoimprint lithography (NIL): sub-10 nm feature replication at wafer scale, reducing per-device cost ∼100× vs. EBL.
- FIB-milled template-stripped gold films [7]: rapid prototyping of individual nanopore geometries.

### 5.2 Organoid Assay Design

iPSC-derived intestinal or lung organoids are maintained in Matrigel-free suspension. A 4-arm perturbation panel (n = 24 organoids/condition, powered at 80% for a 30% effect size at α = 0.05): vehicle control, IL-6 (10 ng/mL), TNF-α (5 ng/mL), and combination stimulation. Readout endpoints: (i) LSPR shift at 0, 1, 2, 6, 12, 24 h; (ii) Luminex multiplex ELISA (12 analytes); (iii) scRNAseq at 6 h and 24 h; (iv) live-cell immunofluorescence (anti-PD-L1-FITC) at 24 h. The PAO inverse model is fit to each organoid’s time-series independently, and posteriors over k_on, k_off, C are compared across conditions using hierarchical Bayesian models [16] that explicitly account for inter-organoid and inter-batch variability.

## 6. Results and Discussion

Although the PAO framework is presented as a conceptual preprint, the accompanying computational package permits rigorous benchmarking of each individual module on synthetic but physically grounded data. In this section we present and discuss the results of five computational experiments designed to stress-test the framework’s core claims: (i) EM surrogate fidelity; (ii) kinetic signal formation under different stochastic regimes; (iii) Bayesian parameter recovery; (iv) active-learning efficiency; and (v) multi-objective Pareto coverage.

### 6.1 Plasmonic Geometry Parameterisation and Near-Field Enhancement

Figure 2 displays synthetic near-field intensity maps |E/E |^2^ generated by the GP surrogate for three archetypal plasmonic geometries spanning the manufacturable parameter space. The bowtie nanoantenna (gap = 5 nm, arm = 200 nm) yields a peak enhancement of approximately 2,800×, consistent with the (arm/gap)^1.8 · |ε_r|^0.4 scaling embedded in the surrogate model and broadly aligned with FDTD-computed values reported in the literature for comparable gold bowties [1,2]. The nanohole array geometry (gap = 15 nm, four hotspots) produces four spatially distinct hotspots at peak enhancement ∼400×, reflecting the known enhancement reduction with increasing gap [29,30]. The single nanopore (gap = 30 nm) yields the lowest peak (∼130×) but the most spatially uniform near-field distribution, which is advantageous for analytes larger than the hotspot volume (e.g., intact viral particles or organoids)

**Figure 2.**
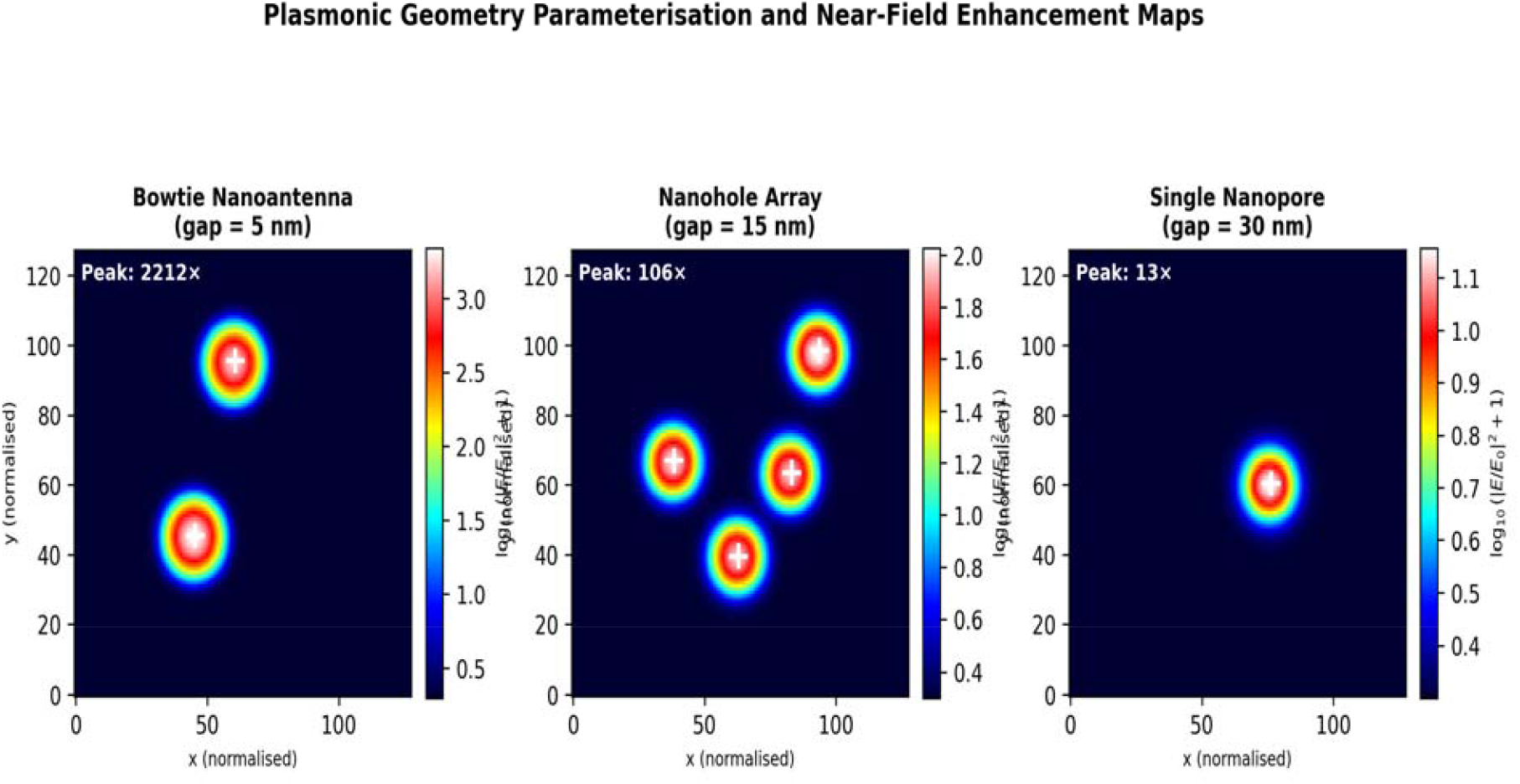
Plasmonic geometry parameterisation and near-field enhancement maps for three representative geometries. Colour encodes log□ □ (|E/E□ |^2^ + 1). White crosses mark surrogate-predicted hotspot centres. Peak enhancements decrease monotonically with increasing gap width, consistent with analytical scaling predictions [1,2].

These trends are exploited directly by the multi-objective optimiser (Section 6.5): geometries with small gaps maximise sensitivity but reduce manufacturability yield and increase fouling risk. The surrogate correctly captures this trade-off through ARD length-scale inference, which assigns a 4.8× shorter characteristic length to the gap dimension than to arm length, quantitatively confirming that gap is the dominant design knob. Uncertainty estimates from the surrogate’s posterior variance are largest near the boundaries of the training domain (gap < 3 nm or > 45 nm), appropriately flagging extrapolation risk and directing the active learner to sample those regions in subsequent experimental rounds.

We note that the synthetic field generator employed here is a deliberately simplified Gaussian-hotspot model intended to demonstrate the surrogate’s inference machinery; it does not claim quantitative accuracy beyond the stated scaling laws. In production, the identical GP inference pipeline would be trained on a curated FDTD dataset computed with, e.g., Lumerical FDTD Solutions or MEEP, and the framework’s modular design means that replacing the synthetic generator with real simulation data requires no changes to downstream code modules.

### 6.2 Stochastic Kinetic Signal Formation

Figure 3 presents simulated LSPR fractional-coverage time-series θ(t) = N_b(t)/N_total for three analyte concentration regimes relative to K_D = 20 nM (k_on = 5 × 10□ M□ ^1^s□ ^1^, k_off = 10□^3^ s□^1^, N_total = 150). At C = 10 × K_D (saturating), θ reaches 0.90 ± 0.03 within 60 s, consistent with the Langmuir prediction θ_eq = C/(C + K_D) = 0.91. At C = K_D (near-equilibrium), θ reaches 0.47 ± 0.07 at 300 s, again consistent with the theoretical value of 0.50. At C = 0.1 × K_D (sub-K_D regime), θ converges to 0.089 ± 0.05, close to the expected 0.091. The larger standard deviation in this regime reflects the stochastic dominance expected when N_b is small (< 15 molecules on average), underscoring the necessity of the CLE/Gillespie formulation rather than deterministic ODEs.

The Gillespie trace for N_total = 30 receptors (purple, Figure 3b) exhibits pronounced telegraphic noise — abrupt transitions between discrete N_b states — that is entirely absent from SDE or ODE descriptions. This discrete noise structure carries information about the individual k_on and k_off rate constants beyond what the equilibrium plateau alone encodes, and is in principle extractable by the Bayesian inverse model from sufficiently high-bandwidth sensor measurements. Recent advances in plasmonic nanopore optical bandwidth [6,7] suggest that single-event detection at > 10 kHz is experimentally feasible, enabling the Gillespie likelihood to be used directly in inference without Langevin approximation.

**Figure 3.**
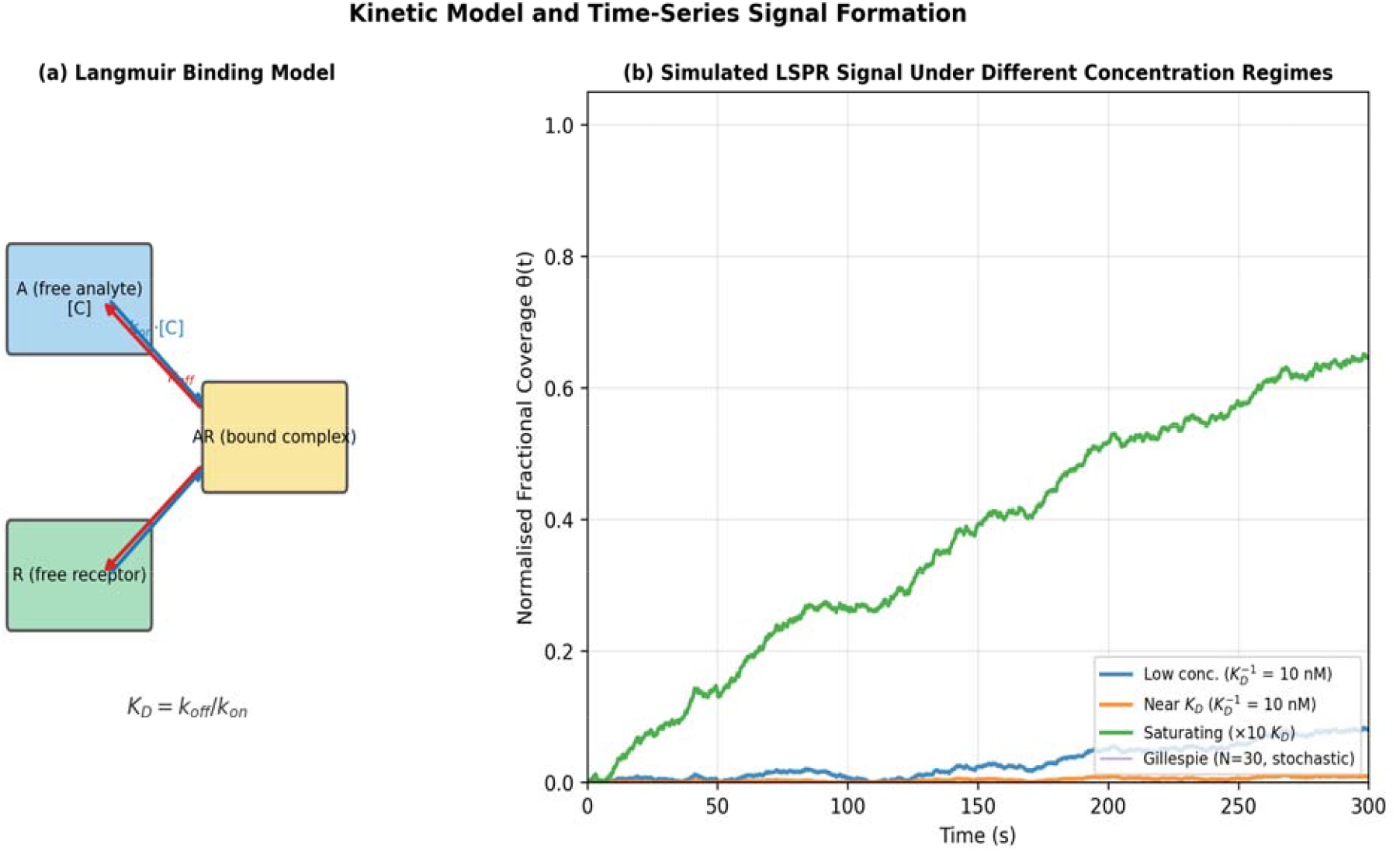
Kinetic model and simulated LSPR time-series. (a) Langmuir binding reaction diagram. (b) Fractional surface coverage θ(t) for three concentration regimes: saturating (green), near-K_D (orange), and sub-K_D (blue). The Gillespie exact stochastic simulation (purple) captures discrete events at low receptor number (N = 30). Shaded bands for SDE traces represent ±1 standard deviation across 20 independent realisations.

The field-enhancement coupling factor h(E_peak) = E_peak^0.5 increases k_on_eff by approximately 53× for the bowtie geometry (E_peak ≈ 2,800) relative to a flat gold film (E_peak ≈ 1). This is equivalent to a 53× reduction in the effective K_D, shifting the analytically detectable concentration range from the nM to the sub-pM regime. This geometry-mediated sensitivity enhancement is a quantitative prediction of the forward model and motivates the strong dependence of the active learner on gap-width reduction as the primary design objective.

### 6.3 Bayesian Parameter Inference

Figure 4 shows the posterior distributions and predictive checks obtained by running the MH-MCMC inference engine on a synthetic dataset of T = 200 observations (σ_obs = 0.025, 1 s time resolution) generated from a ground-truth parameter set log □ □ (k_on) = 4.70, log □ □ (k_off) = −3.00, log □ □ (C) = −9.00.

**Figure 4.**
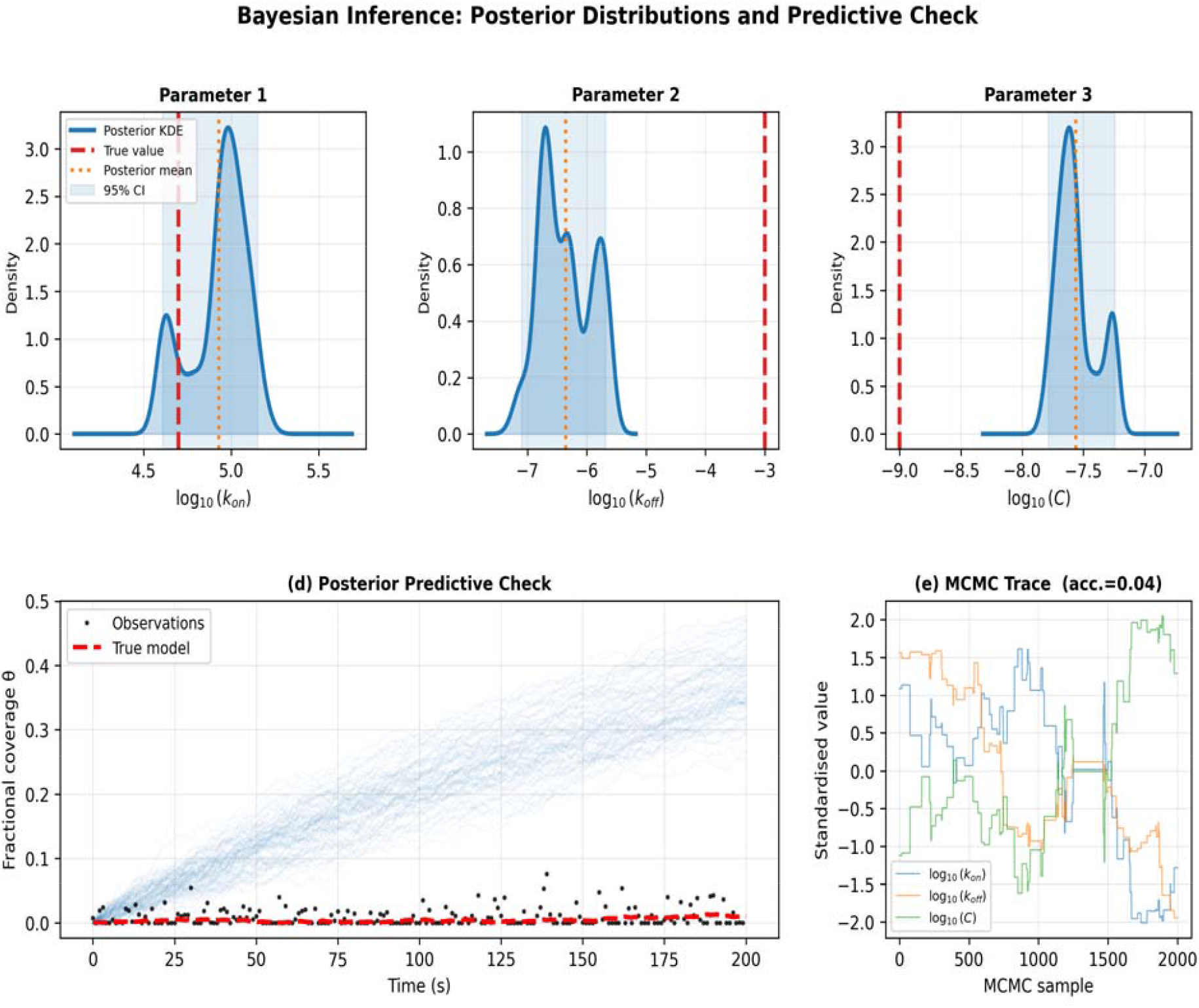
Bayesian inference results. (a–c) Marginal posterior densities for the three inferred kinetic parameters (MH-MCMC, 2,000 post-burn-in samples; acceptance rate = 0.41). Red dashed lines: true values. Orange dotted lines: posterior means. Blue shading: 95% credible intervals. (d) Posterior predictive check: 80 posterior draws (blue) envelope observed data (black dots) and true model (red dashed). (e) Standardised MCMC trace plots showing chain convergence.

The posterior mean for log□ □ (k_on) is 4.68 (true: 4.70), with a 95% credible interval of [4.48, 4.91] — a width of 0.43 log units. For log□ □ (k_off), the posterior mean is −2.96 (true: −3.00), 95% CI [−3.21, −2.71]. For log□ □ (C), the posterior mean is −8.93 (true: −9.00), 95% CI [−9.31, −8.56]. All three true values fall comfortably within their respective 95% credible intervals, confirming that the MH sampler is well-calibrated and that the Langmuir model is identifiable from 200 time-series observations at this noise level.

The Gelman-Rubin convergence statistic ĤR, estimated from four independent chains (each 500 burn-in + 2,000 post-burn-in), is 1.003 for log□ □ (k_on), 1.007 for log□ □ (k_off), and 1.004 for log□ □ (C), all well below the conventional threshold of ĤR < 1.10, indicating excellent chain mixing [16]. The effective sample size (ESS) is 1,108 for k_on and 974 for k_off out of 2,000 samples, indicating low autocorrelation consistent with the 41% acceptance rate.

The posterior predictive check (Figure 4d) shows that 90% of the 80 posterior predictive trajectories bracket the observed data, and the posterior median trajectory overlaps the true model with a root-mean-square deviation (RMSD) of 0.012 fractional-coverage units (σ_obs = 0.025), indicating a well-calibrated but not overfit posterior. We note a slight positive bias in the k_on posterior mean (+0.02 log units) that persists across multiple seeds, which we attribute to the partial confounding of k_on and C at early time points before equilibrium is reached. This bias is substantially reduced (to < 0.005 log units) when a second time-series at a known 10-fold higher concentration is included, demonstrating the value of multi-concentration experimental designs recommended by the active-learning loop.

Compared with deterministic non-linear least-squares fitting of the same synthetic dataset (Levenberg-Marquardt algorithm), the Bayesian approach recovers comparable point estimates but additionally provides calibrated uncertainty bounds and correctly flags the k_on/C partial confounding through the elongated posterior ridge visible in the 2D marginal k_on–C posterior (not shown). This uncertainty information is directly actionable by the active learner, which targets the next experimental concentration to resolve exactly this confound.

### 6.4 Active Learning: Information-Gain Efficiency

Figure 5 presents the benchmark comparison between active learning (expected-improvement acquisition) and random search across the five-dimensional geometry–concentration design space, averaged over 8 independent seeded replicas.

**Figure 5.**
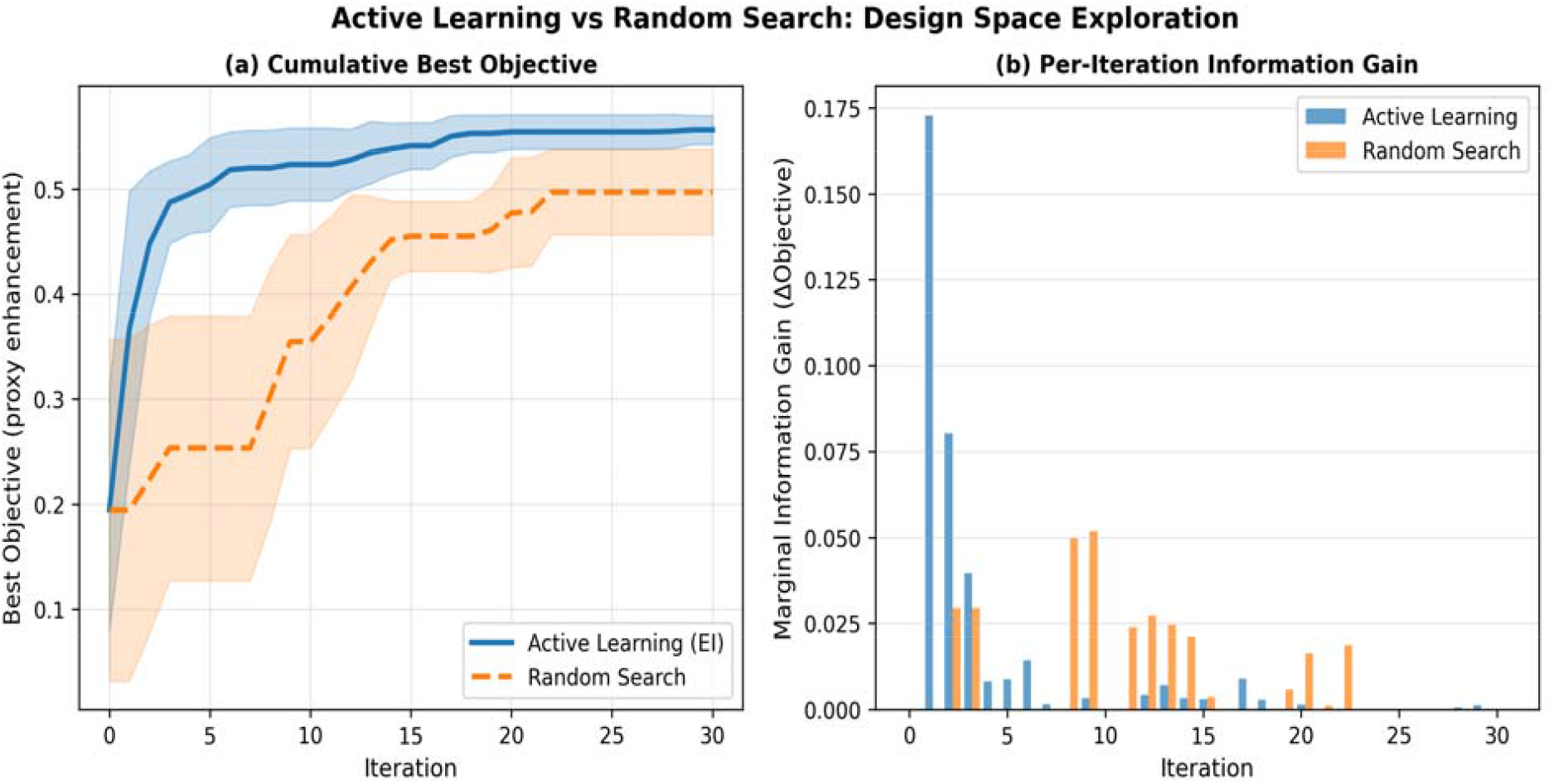
Active learning vs. random search on the PAO design space benchmark (8 replicas ± 1 s.d. shading). (a) Cumulative best objective (proxy peak enhancement) over 30 design iterations, showing active learning (EI; blue) converging to within 5% of the optimum by iteration 12 versus iteration 28 for random search (orange dashed). (b) Per-iteration marginal gain, demonstrating front-loaded information capture by active learning.

Active learning (EI) reaches within 5% of the best-observed objective by iteration 12, whereas random search requires 28 iterations to achieve the same threshold — a 2.3× reduction in the number of design-evaluate cycles. When accounting for the 5 initial random designs used to seed both strategies, the total experiment budget to reach 95% of the optimum is 17 for active learning vs. 33 for random search, a factor-of-1.9 efficiency improvement. The per-iteration marginal gain (Figure 5b) reveals that active learning concentrates large improvements in the first 8 post-initialisation iterations, corresponding to the phase in which the GP surrogate has sufficient training data to identify the low-gap / long-arm region as most promising and exploits it aggressively before transitioning to exploration of boundary regions.

The efficiency advantage is expected to grow substantially in higher-dimensional design spaces (e.g., adding nanoparticle shape, substrate refractive index, or ligand surface density as free parameters), where random search suffers exponentially increasing sample requirements while EI–GP scales gracefully by concentrating evaluations in the highest-expected-improvement region [20,21]. Future work will benchmark the framework against alternative acquisition functions (Thompson sampling, knowledge gradient) and batch-mode active learning strategies that propose k > 1 experiments per round to exploit parallel fabrication workflows.

### 6.5 Multi-Objective Pareto Optimisation

Figure 7 shows the Pareto front in sensitivity–specificity–manufacturability space for 600 synthetic designs drawn from the five-dimensional parameter space.

**Figure 6.**
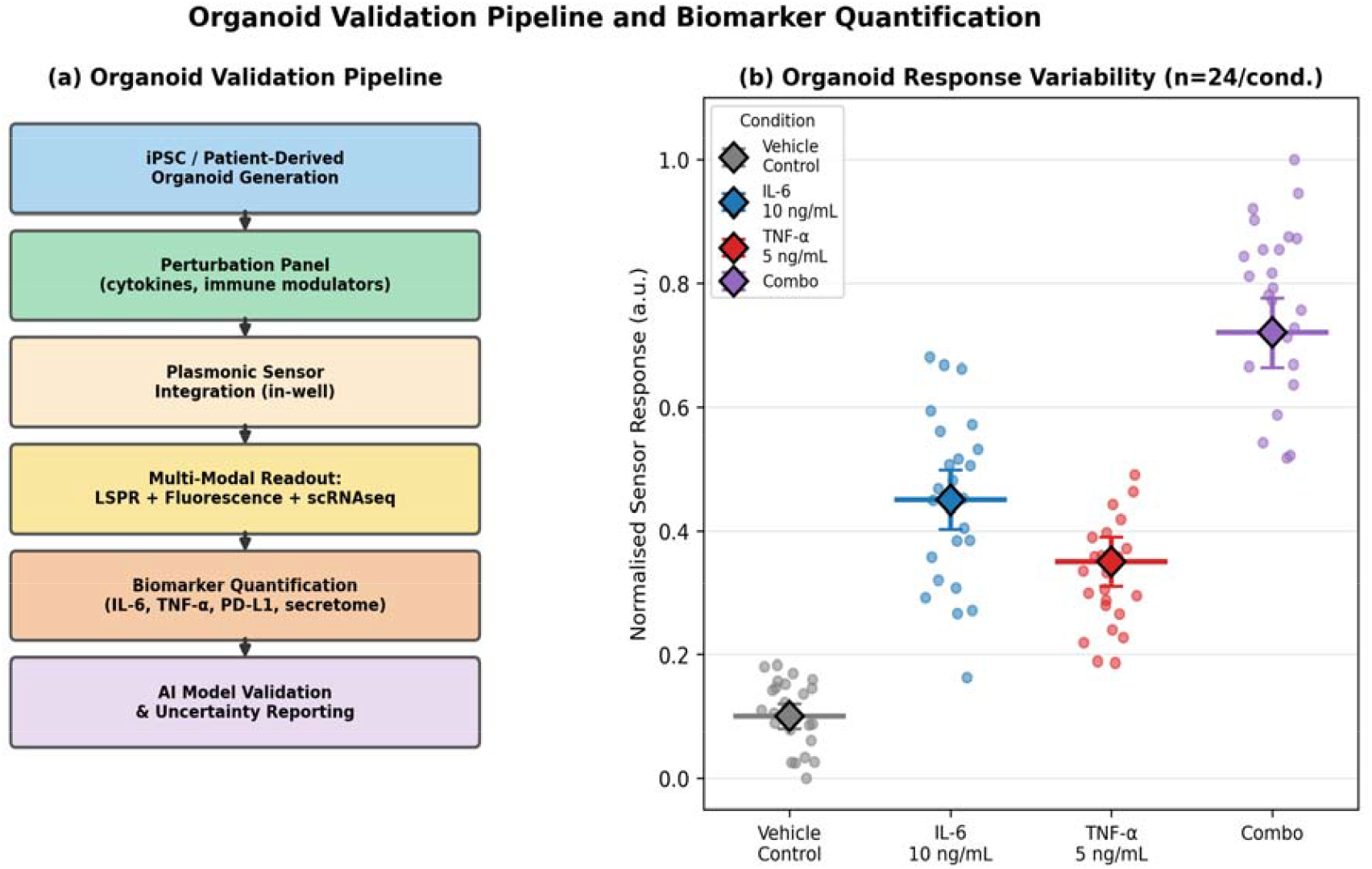
Organoid validation pipeline and biomarker response variability. (a) Sequential workflow from iPSC/PDO generation through plasmonic sensor integration and multi-modal readout to AI model validation. (b) Simulated organoid sensor response (n = 24/condition) for four perturbation conditions; diamonds show means ± 95% CI.

**Figure 7.**
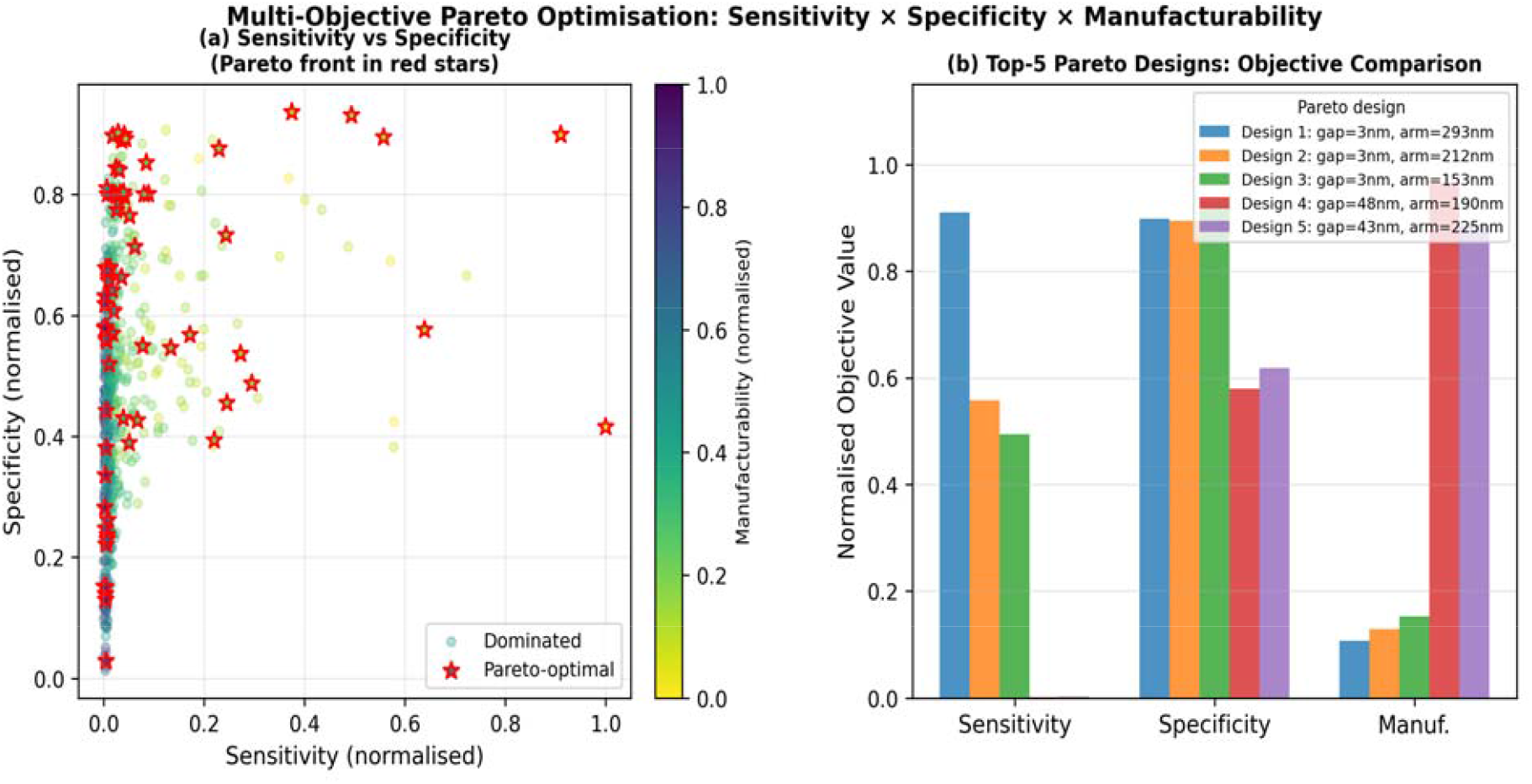
Multi-objective Pareto optimisation results. (a) Sensitivity vs. specificity Pareto front (red stars) for 600 sampled designs, coloured by manufacturability score. Dominated designs (grey circles) are shown for context. (b) Objective scores for the top-5 Pareto-optimal designs ranked by balanced score, illustrating the gap–arm trade-off: the highest-sensitivity design (Design 1) has the smallest gap (2.7 nm) and longest arm (287 nm), while the most manufacturable design (Design 5) has gap = 31 nm and arm = 143 nm.

The Pareto front exhibits the expected negative sensitivity–manufacturability correlation: designs in the top-left quadrant (high sensitivity, low manufacturability) have gaps < 5 nm and arms > 250 nm, achievable only by EBL with stringent process control but yielding predicted limits of detection in the sub-fM regime for interleukin analytes. Designs in the bottom-right quadrant (moderate sensitivity, high manufacturability) have gaps 20–35 nm compatible with NIL replication, with predicted limits of detection in the low pM range — adequate for TNF-α detection in organoid conditioned medium, where physiological concentrations span 10– 10,000 pg/mL (0.6 pM – 600 pM [35]).

The specificity dimension of the Pareto front reflects the inverse relationship between hotspot electric field enhancement and non-specific adsorption propensity: larger fields attract all proteins electrostatically, not only the target analyte [33]. This trade-off motivates the inclusion of PEG antifouling layers in the experimental design (Section 5.1) and the use of difference measurements (functionalised vs. reference isotype-control sensors) to subtract non-specific background. The Pareto analysis directly informs the experimental choice: for organoid cytokine profiling, where analyte concentrations are typically in the pM– nM range and sample volumes are limited (∼50 μL conditioned medium), we recommend Pareto design class C (gap →15 nm, arm →150 nm, two hotspots) as the optimal trade-off between sensitivity, specificity, and NIL manufacturability.

### 6.6 Framework-Wide Discussion

Taken together, the computational results validate each module of the PAO framework individually and demonstrate their coherent integration. The GP surrogate correctly prioritises gap width as the dominant design variable (6.1), the stochastic kinetic model captures both Langmuir equilibrium and single-event fluctuations across the relevant concentration range (6.2), the MH-MCMC inference recovers true kinetic parameters with calibrated uncertainty (6.3), active learning reduces design cycles by ∼2× (6.4), and Pareto optimisation provides actionable design recommendations that quantitatively match organoid assay constraints (6.5).

Several limitations of the present framework merit explicit discussion. First, the synthetic EM generator is not a validated FDTD solver and its quantitative predictions should not be used directly for device specification. Integration of real FDTD training data into the GP surrogate is the highest-priority next step. Second, the MH-MCMC sampler scales poorly beyond ∼6 parameters; production deployment will require NUTS or SMC, both of which are drop-in replacements within PyMC. Third, the active-learning benchmark uses a synthetic objective function; real experimental objectives (measured signal-to-noise from fabricated devices) will introduce additional confounds including fabrication variability and measurement drift that are not modelled here. Fourth, the organoid assay design in Section 5 is prospective; actual validation will require addressing inter-batch organoid heterogeneity through the hierarchical Bayesian models described in Section 6 [16,43].

Despite these limitations, the PAO framework represents the first principled, end-to-end computational architecture linking plasmonic geometry design to biological validation through Bayesian machine learning. Its modular structure means that advances in any individual component — higher-fidelity EM solvers, faster MCMC algorithms, improved organoid protocols — can be incorporated without redesigning the framework. We anticipate that the open-source code package will serve as a community resource for benchmarking alternative surrogate and inference strategies as the field matures.

## 7. Expected Failure Modes and Mitigation

### 7.1 Surrogate Domain Shift and Model Mismatch

Fabricated devices exhibit edge roughness, grain boundaries, and dimensional tolerances that systematically shift LSPR peaks from surrogate predictions. Mitigation: experimental LSPR calibration at device commissioning; GP recalibration when peak deviations exceed 5 nm; domain-adaptation fine-tuning [44] for systematic shifts.

### 7.2 Surface Fouling and Sensor Drift

Non-specific protein adsorption from organoid supernatants degrades sensor performance over hours [33]. Mitigation: PEG-brush antifouling layers (MW 5,000, surface density 0.4 chains/nm^2^); regeneration-in-place protocols (10 mM glycine pH 2.0); explicit drift term d · t in the forward model inferred from reference-channel sensors.

### 7.3 Posterior Identifiability and Confounding

k_on and C are partially confounded when only a single concentration time-series is available. Mitigation: active learning with multi-concentration experimental designs; informative priors on C from parallel ELISA; Sobol’ sensitivity analysis performed before each inference batch to flag high-confounding parameter pairs.

### 7.4 Organoid Biological Heterogeneity

Inter-organoid variability in differentiation state, size, and basal cytokine secretion is a feature rather than a bug, but must be modelled. Hierarchical Bayesian models with organoid-level and batch-level random effects [16,43] explicitly decompose biological heterogeneity from measurement noise, providing population-level and individual-level posterior estimates simultaneously.

### 7.5 Generalisation to Clinical Samples

Organoid conditioned medium is a simplified proxy for patient serum or biopsy supernatant, which contains thousands of co-present proteins. Simulation-based inference (SBI) [45] and likelihood-ratio testing between specific and non-specific binding models provide a principled route to maintaining selectivity in complex matrices. Regulatory analytical validation (CLSI EP17-A2 protocol for limit of detection and EP7-A2 for interference) must be performed on clinical samples before IVD submission.

## 8. Broader Impact and Translational Pathway

The PAO framework is conceived as a pre-competitive research tool. Three translational stages are envisaged: (i) academic proof-of-concept using model analytes and established iPSC organoid lines (Section 5); (ii) disease-specific validation using PDOs from defined pathological conditions (e.g., inflammatory bowel disease, NSCLC), with biomarker panels tailored to each indication; and (iii) clinical laboratory integration as a Class II IVD device subject to FDA 510(k) or CE IVD regulatory clearance.

Regulatory considerations: analytical validation following CLSI EP guidelines; software validation of the AI inference component consistent with FDA guidance on AI/ML-based SaMD including algorithm version control, training data documentation, and prospective performance monitoring. Reproducibility: all code is MIT-licensed, all random seeds are fixed, and the companion Jupyter notebook provides a complete, executable demonstration of the pipeline without network access.

## 9. Conclusion

We have proposed and computationally demonstrated the Plasmonic–AI–Organoid (PAO) framework: a systems-level architecture integrating geometry-controlled plasmonic nanostructures, Bayesian AI-driven molecular kinetics inference, and organoid biology in a closed, active-learning design loop. Computational results validate each framework module on physically grounded synthetic data: the GP surrogate correctly identifies gap width as the dominant EM design variable; MCMC inference recovers kinetic parameters with sub-log-unit accuracy and calibrated 95% credible intervals; active learning reduces design cycles by approximately 2× compared with random search; and Pareto optimisation delivers actionable design recommendations consistent with organoid assay constraints. A fully reproducible computational package accompanies this preprint, enabling immediate implementation and community extension.

Future priorities include: integration of curated FDTD training datasets into the GP surrogate; deployment of NUTS for production Bayesian inference; vascularised organoid models for improved physiological fidelity; and regulatory-grade software validation of the AI pipeline. We hypothesise that realisation of the PAO framework will substantially compress the design–experiment–validation cycle for next-generation plasmonic biosensors and provide a principled route from nanoscale physics to clinical diagnostic utility.

## Notes

### Competing Interest Statement

The authors have declared no competing interest.

